# Machine Learning-Driven Multiplexed Biomarker Detection with Polymer-Enhanced Electrochemical Sensors

**DOI:** 10.64898/2026.05.04.722575

**Authors:** Adrian L. M. Duesselberg, Ines C. Weber, Yann Zosso, Parastoo Salah, Zhenan Bao

## Abstract

Biomarkers in sweat and saliva offer a promising avenue for non-invasive health monitoring. Electrochemical sensors have the potential to measure such biomarkers simultaneously. However, they are limited in discriminating individual biomarkers in mixtures, as redox potentials often overlap, resulting in current signatures that cannot be deconvoluted.

This study focuses on differentiating biomarkers using orthogonal sensing materials combined with machine learning. We introduce a flexible electrochemical sensor array comprising carbon flower electrodes modified with poly(vinylidene fluoride) (PVDF) or poly(4-vinylpyridine) (P4VP) for the detection of estradiol (E2), ascorbic acid (AA), serotonin (5-HT), and melatonin (Mel). The two polymers act by altering the redox potential and current response of each biomarker, thereby enhancing signal diversity and enabling peak separation. Using multi-output regression models on 450 single and mixture measurements, the array accurately predicts concentrations (*R*^2^ = 0.95) over a wide dynamic range spanning nanomolar to micromolar levels. Polymer-resolved analysis reveals that PVDF-modifications enhance E2 and Mel detection, while P4VP-modifications improve AA and 5-HT quantification, highlighting the benefit of complementary orthogonal sensing electrodes. This finding is further supported by feature attribution analysis, which shows that the machine learning model relies on polymer-specific electrochemical signatures, directly linking improved performance to distinct polymer-analyte interactions. Overall, these results demonstrate that combining polymer-modified orthogonal electrodes with machine learning enables accurate, multiplexed sensing in complex mixtures, advancing selective detection strategies for future sensor platforms.

## Introduction

Human sweat and saliva contain numerous biomarkers that reflect biological processes within the body. Differences in biomarker concentrations provide insights into healthy and diseased states, making them attractive for non-invasive disease diagnostics and continuous health monitoring for personalized and preventive healthcare.^1^ For example, estradiol (E2) is a hormone essential in female health^2^ and affects major organs, including bones^3^ and the brain.^4^ Serotonin (5-HT) regulates mood and is linked to neurological disorders, such as depression and post-traumatic stress disorder,^5^ while peripheral serotonin is related to metabolic health.^6^ Ascorbic acid (AA) is an essential antioxidant and nutritional indicator.^7^ Melatonin (Mel) influences circadian rhythms and sleep patterns.^8^

For this reason, wearable sensors have attracted increasing interest.^9–11^ Carbon-based electrochemical sensors are among the most widely studied types, as they are low-cost, have high potential stability, and are reversible. Various electrochemical neurotransmitter and hormone sensors have been reported in recent years with high sensitivity using strategies including metal (oxide) doping,^12,13^ high surface area materials,^14^ and conductive polymers. ^15^ However, accurately detecting and quantifying specific biomarkers remains a significant challenge due to their co-occurrence and overlapping redox potentials, resulting in overlapping current signatures that cannot be deconvoluted. In addition, sensors often do not meet the mechanical requirements for soft, on-skin applications due to their rigidity, for example, modified glassy carbon electrodes (GCEs), or toxicity (i.e., carbon nanotubes).

Sensor arrays offer a solution towards multiplexed sensing. ^16^ In principle, each electrode provides an individual signal pattern, generating diverse and distinguishable signal sets that enable identification and quantification of individual components. While sensor arrays provide a viable route towards selectivity, their signal readout, processing, and interpretation remain complex, particularly when analyte responses overlap strongly. Machine learning (ML) has therefore emerged as a powerful complementary tool to extract multivariate information from electrochemical signals and enable multiplexed quantification beyond conventional peak-based analysis. Recent studies illustrate this trend: overlapping dopamine (DA) and 5-HT signals were distinguished using artificial neural networks (ANNs), achieving *R*^2^ *>* 0.97 and root mean squared error (*RMSE*) *<* 0.1 µM;^17^ linear regression models were applied to cyclic voltammetry data (*n* = 45) to distinguish glucose and insulin, yielding *R*^2^ *>* 0.98 with 4 − 6% error; ^18^ multiplexed detection was demonstrated on engineered sensor surfaces with supervised ML models (support vector machine (SVM), ANN) to distinguish tyrosine and uric acid in sweat and saliva, reaching *R*^2^ = 0.85 − 0.95;^19^ and SVM, ANN, and GLM-NET regression were used to detect multiple toxicants in water, reporting *RMSE* ≈ 0.24 mg *L*^−1^ across 23 sensors.^20^ Despite these advances, many prior works are constrained by limited dataset sizes, limited analyte diversity, and a lack of distinctly different electrodes. Such limitations can hinder model generalization and increase the risk of overfitting when applied to realistic sensing scenarios. ^21^ This emphasizes the need for larger, more realistic, and multiplexed datasets^22,23^ alongside improvements in sensor design and electrode specificity, model robustness, and standardized protocols for deploying ML in practical sensing applications.^24–26^

In one of our recent studies, we demonstrated that introducing polymer modifications with poly(4-vinylpyridine) (P4VP) and poly(vinylidene fluoride) (PVDF) induces polymer-specific peak potential shifts due to different adsorption and diffusion characteristics of individual biomarkers with these polymers.^27^ We further showed that these polymer modifications work well with nanostructured carbon flowers to achieve high sensitivity. ^28^ In this work, a sensor array based on polymer-modified carbon flowers is designed to detect the biomarkers E2, AA, 5-HT, and Mel (**Fig. 1a**). Sensor arrays are fabricated by spray-coating carbon flower inks containing the respective PVDF or P4VP polymers onto stretchable polystyrene-block-poly(ethylene-ran-butylene)-block-polystyrene (SEBS) substrates. We collected a comprehensive dataset of 450 measurements comprising 1-, 2-, 3-, and 4-analyte mixtures to capture realistic mixed-analyte conditions. This platform is combined with a structured ML framework to identify analyte concentrations from multiplexed voltammetric signals. The framework integrates linear, kernel-based, ensemble, and neural network models to examine how algorithmic flexibility, preprocessing transformations, and feature representations affect predictive performance and robustness. Interpretability is ensured through SHapley Additive exPlanations (SHAP)-based analysis, ^29^ enabling identification of signal regions and polymer-specific interactions most relevant to prediction. Together, these strategies address longstanding challenges in selective electrochemical detection of biomarkers in complex mixtures and demonstrate the potential of polymer-orthogonal sensing combined with ML for wearable and point-of-care applications.

**Figure 1:**
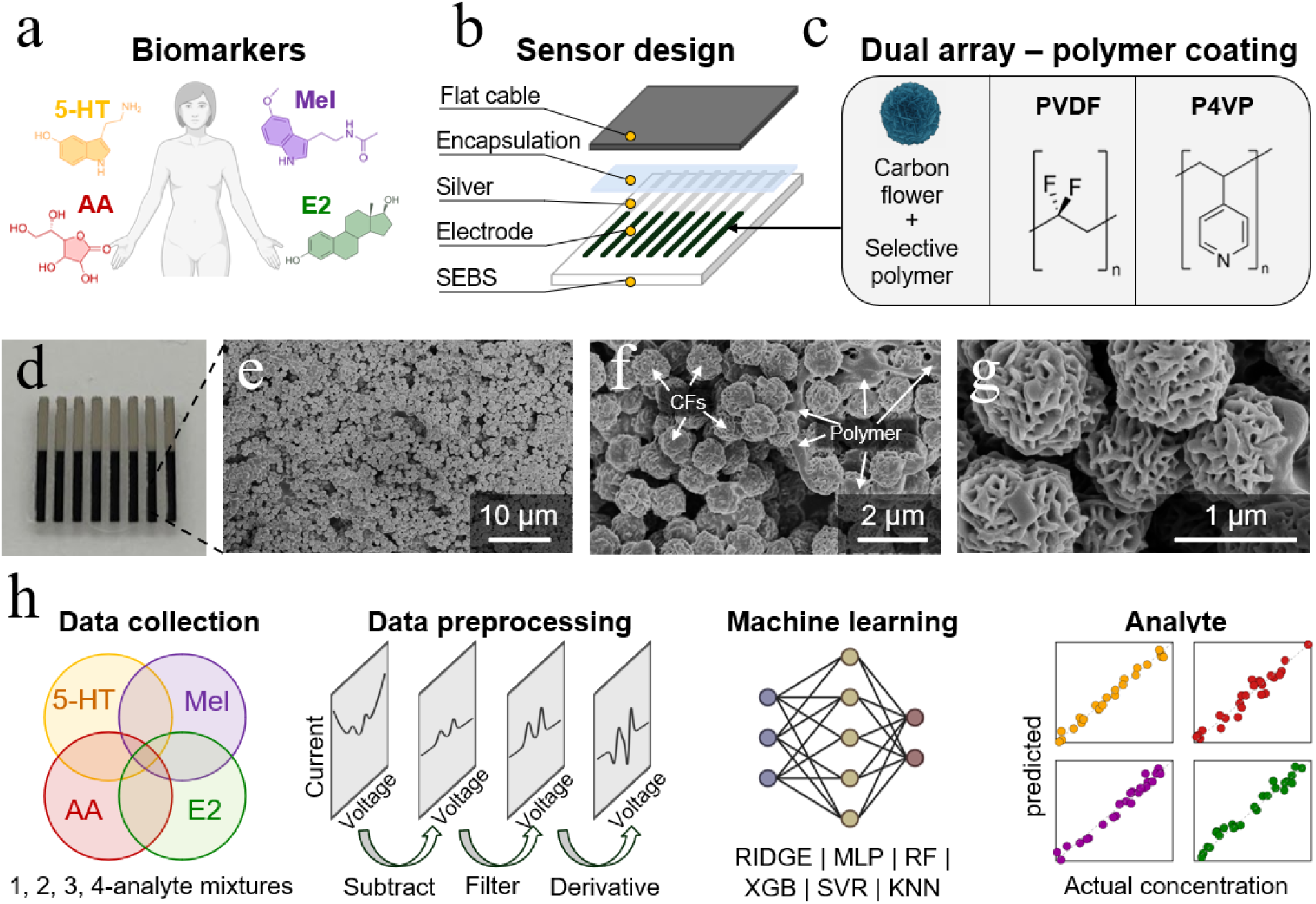
— Concept. **(a)** Target non-invasive biomarkers: estradiol (E2), serotonin (5-HT), ascorbic acid (AA), and melatonin (Mel). **(b)** Schematic of sensor design. **(c)** Dual sensor array: each sensor consists of 8 electrodes made of conductive carbon flowers coated with selective polymers — Type 1: PVDF-CF, Type 2: P4VP-CF. **(d)** Photograph of the electrochemical sensor. **(e-g)** SEM image of an electrode and zoomed-in views. **(h)** Data acquisition and ML workflow.

## Experimental Section

### Sensor Fabrication and Characterization

The sensor fabrication followed the protocols from our previous works. ^28,30,31^ Carbon flowers were synthesized from acrylonitrile precursors and subsequently carbonized at 1500 °C under nitrogen flow. The as-prepared carbon flowers (45 mg) were dispersed in chloroform (100 mL) containing the respective polymers (5 mg), poly(4-vinylpyridine) (P4VP, *M*_*w*_ = 60,000 Da, Sigma-Aldrich) or poly(vinylidene fluoride) (PVDF, *M*_*w*_ = 534,000 Da, Sigma Aldrich), using ultrasonication to obtain a homogeneous ink. The carbon flower inks were then spray-coated onto stretchable SEBS (Asahi Kasei Tuftec H1052, Tokyo, Japan) sub-strates featuring pre-defined stretchable silver current collector patterns, using a metal mask to define the electrode geometry. Electrical connections were established using conductive *z*-axis tape to interface the carbon flower electrodes with a flat cable connector. Each sensor comprised eight working electrodes mixed with the same polymer formulation (PVDF or P4VP); only one electrode was used for electrochemical testing. In total, 5 P4VP-modified sensors (ID: MC302, MC312, MC327, MC329, MC330) and 3 PVDF-modified sensors (ID: MC313, MC320, MC325) were fabricated and included in this study. Scanning electron microscopy was conducted using an FEI Magellan 400 XHR instrument, operated at an accelerating voltage of 5 kV and a beam current of 100 pA.

### Electrochemical Measurements

Electrochemical measurements were performed using a CH Instruments 760E potentiostat coupled with a CHI200 picoamp booster and Faraday cage to reduce electrical noise. Square wave voltammetry (SWV) was used for analyte detection in phosphate-buffered saline (PBS) using the following parameters: 0.025 V pulse amplitude, 5 Hz frequency, and 0.004 V step potential across a 0–0.8 V range (resulting in 200 current values per trace). A standard three-electrode setup was used, consisting of a platinum wire auxiliary electrode, an Ag/AgCl (3 M KCl) reference electrode, and the respective carbon flower working electrode modified with either PVDF or P4VP. The CHI setup enabled simultaneous measurements of two electrodes, resulting in two similar current responses per scan. To avoid redundant information in subsequent modeling, only the first current trace was used for preprocessing and ML analysis. Experiments were conducted at ambient temperature in 1*×* PBS (pH 7.2). At least three blank scans were collected at the start of the measurement sequence to capture the plain baseline PBS response for subsequent data preprocessing. 17*β*-estradiol (*>* 0.97, TCI Chemicals) and melatonin stock solutions were prepared in ethanol, whereas serotonin hydrochloride (≥ 98, Sigma Aldrich), L-ascorbic acid (≥ 0.98, Sigma Aldrich), uric acid (≥ 98, AmBeed), dopamine hydrochloride (≥ 0.98, Sigma Aldrich), norepinephrine (≥ 0.98, Sigma Aldrich), and epinephrine (≥ 0.99, Sigma Aldrich) were dissolved directly in PBS. Analyte solutions were freshly prepared and introduced stepwise into nitrogen-purged PBS at specified concentrations. Mixing was accomplished through nitrogen bubbling. Typically, three consecutive measurements were performed at each concentration step, with a uniform waiting time of 60 s; for later analysis, the average of these scans was used. Prior to each measurement, sensors were rinsed with deionized water.

### Dataset Design and Concentration Ranges

For each polymer coating, the dataset was designed to include a diverse set of mixture complexities: 10 single samples per biomarker (80 total), 15 samples for each of six biomarker pairs (180 total), 15 samples for each of four triple combinations (120 total), 15 samples for each quadruple (30 total), and 40 additional random samples. In total, this resulted in 450 samples. Detailed measurement and data parameters are provided in **Table S1**, and a full dataset summary across polymer coating and individual sensors is shown in **Table S2**. The concentration ranges were selected based on reported physiological concentration levels for ascorbic acid and serotonin in relevant biofluids (see **Table S3**); for estradiol and melatonin, concentration levels higher than physiological levels were chosen due to the sensitivity limitations of the sensors. For PVDF-modified sensors, concentrations ranged from 10–1000 nM for E2, 5-HT, and Mel and 1–10 *µ*M for AA. For P4VP-modified sensors, concentrations ranged from 100–3000 nM for E2, 5-HT, and Mel and 6–20 *µ*M for AA.

### Data Preprocessing, Machine Learning, and Performance Evaluation

Raw SWV scans (**Fig. S1a**, 200 data points per trace) were preprocessed to reduce measurement noise and enhance electrochemical features. First, up to three blank PBS scans were averaged and subtracted from the analyte signals, followed by Savitzky–Golay (SG) filtering (polynomial fit of degree 3 over a 9-point window) to suppress high-frequency noise while retaining peak shapes and overall signal morphology (**Fig. S1b**). The potentiostat used in this study (CHI750E) produces very little noise, with the high-frequency component representing only ∼3.5% of the total signal variance (SNR ≈ 276). SG smoothing provides approximately 2*×* noise reduction in the raw signal and 2.8*×* in the derivative domain. These steps ensured that models were trained on signals reflecting true electrochemical activity rather than artifacts from instrumentation or baseline drift.

To address baseline variability and overlapping peaks, three signal transformation strategies (**Fig. S1c**) were evaluated: min-centering for baseline alignment, linear baseline correction by fitting and subtracting a first-degree polynomial using the edge regions of the voltammogram (10 points at the start and end), and first-derivative transformation (*dI/dV*) computed via numerical gradient (`numpy.gradient`), which enhances peak contrast and introduces secondary extrema to improve analyte separability. If not noted explicitly otherwise, first-derivative transformation was used, as it yielded the best performance.

For ML analysis, the preprocessed scans were used directly, allowing the models to learn from the complete voltammetric signal without assuming a fixed number or position of peaks. In addition to the electrochemical features, two categorical variables were included: the sensor identifier (ID number) and the polymer coating type.

The modeling task was formulated as supervised multi-output regression to predict concentrations of the four biomarkers (E2, AA, Mel, and 5-HT). To prevent data leakage from repeated measurements, samples were grouped by sample ID during splitting. For each random seed (0, 42, 1337, 2023, 9999), an independent held-out test set was generated using GroupShuffleSplit (90/10). The remaining 90% of the data was used for model development with 5-fold GroupKFold cross-validation. Training and validation metrics were computed as the mean across the five cross-validation folds for each seed, while test metrics were evaluated on the corresponding held-out test set. All reported performance values are given as mean *±* standard deviation across the five random seeds. Numerical features were standardized to zero mean and unit variance using a `StandardScaler`, fitted exclusively on the training data, and subsequently applied to the validation and test sets, while categorical features were one-hot encoded. Six regression algorithms representing diverse learning paradigms were implemented to explore how model complexity and inductive bias influence prediction from multiplexed voltammetric signals: Ridge regression (serving as a baseline for linear modeling with regularization), random forest and XGBoost (tree-based ensembles, suited for non-linear tabular data), *k*-nearest neighbours (non-parametric local model), support vector regression (kernel-based method for non-linear signal relationships), and multilayer perceptron (MLP, a small neural network capable of capturing complex non-linearities). Hyperparameters were tuned using grid search cross-validation and were tuned inside the cross-validation loop; the held-out test set was not used for model selection. Parameter ranges were guided by prior biosensing ML literature^22,23,32^ and refined iteratively.

False non-zero predictions, where the model predicts non-zero concentrations in the absence of an analyte, were mitigated using a zero-aware two-stage architecture for the representative model shown in Fig. 3. In this approach, each analyte was first classified as present or absent using multi-output binary classification (labels: *y >* 0), and concentrations were then predicted using multi-output regression. At inference time, regression outputs were gated by the classifier decision and set to zero when an analyte was predicted absent. This suppresses “phantom positives” while retaining the regression performance on samples where the analyte is present. Negative concentration predictions were clipped to zero prior to evaluation to enforce physical plausibility, which can yield exact 100% relative errors, when non-zero ground-truth samples are predicted as zero.

Predictive accuracy was assessed using complementary metrics. The coefficient of determination (*R*^2^) quantified variance explained, while root mean square error (RMSE) emphasized large residuals, which are particularly important in biosensing applications where outliers can represent sensor artifacts. Mean absolute error (MAE) provided a more interpretable measure of average prediction error across analytes. Training and validation scores were retained to assess potential overfitting, and predicted-versus-true plots were used to visualize systematic deviations. Residual distributions were additionally inspected to identify potential systematic biases across concentration ranges.

SHAP feature importance was computed using KernelExplainer with 100 explanation samples and 100 background samples per fold (randomly sampled from a pool of 200 samples in the training set). SHAP values were computed for the representative split (seed = 42) and averaged across the five cross-validation folds. To assess stability, we verified that feature rankings remain stable across different background sample sizes (**Fig. S2**).

## Results and Discussion

### Sensor Array

Wearable sensor arrays require skin conformability, low-cost production, simple fabrication, and high-throughput capability. These requirements are addressed in our fabrication procedure. The sensor design is depicted in **Fig. 1b** and consists of a stretchable SEBS substrate, conductive stretchable silver interconnects, orthogonal carbon-based electrodes, an SEBS encapsulation layer, and a flat cable connector. The sensing electrodes are fabricated by spray-coating electrode inks comprising carbon flowers with either PVDF or P4VP (**Fig. 1c**). This simple and robust fabrication technique is well-suited for sensor arrays and can be readily adapted and scaled up to incorporate additional polymer modifications.

A photograph of the as-prepared PVDF-carbon flower (PVDF-CF) sensing electrodes before encapsulation and flat cable connection is shown in **Fig. 1d**. The SEBS substrate is flexible and skin-conformable,^28^ ideal for wearable applications. A representative SEM image of the sensing electrode is shown in **Fig. 1e**, with a magnified view in **Fig. 1f**. The carbon flowers have a diameter of approximately 1 µm (**Fig. 1g**) and are connected by polymer binder. Carbon flowers are used due to their beneficial properties in electrochemical sensing, which have been attributed to their unique morphology, defect-rich structure, and high conductivity. ^28^ Importantly, while the polymer binder does not cover all carbon flowers homogeneously, we have previously demonstrated that the polymer effect on electrochemical performance is retained.^27^

The data collection and ML workflow are shown in **Fig. 1h**. A comprehensive dataset of 450 samples was collected, comprising single-analyte, 2-, 3-, and 4-analyte mixtures. Data preprocessing included blank subtraction, filtering, and derivative transformation (**Fig. S1**). ML algorithms, including Ridge regression, multilayer perceptron (MLP), random forest (RF), XGBoost (XGB), support vector regression (SVR), and k-nearest neighbors (KNN), were trained on the preprocessed data, and the best-performing algorithm was used to predict analyte concentrations. Overall, this modular platform architecture combined with widely available ML methods enables straightforward integration of additional electrode types to diversify the feature set, thereby facilitating distinction of a broader range of molecules in complex mixtures.

### Electrode-specific Electrochemical Detection

Representative voltammograms illustrating the effect of the polymer modifications are shown in **Fig. 2** (raw data **Fig. S3**). To emphasize the oxidation potential shift, we show the normalized voltammograms for each biomarker in **Fig. 2a**. Each polymer–biomarker pair exhibits a characteristic oxidation potential, reflecting the polymer-dependent interaction. For example, E2 shows a clear oxidation peak at 0.50 V on PVDF-CF. In contrast, P4VP-CF electrodes detect estradiol at 0.64 V. When testing different concentrations, the peak current varies while the oxidation potential remains unchanged. This significant 140 mV shift in oxidation potential is characteristic of the polymer coating and crucial for achieving selective detection. This likely relates to altered diffusion and adsorption characteristics.^27^

**Figure 2:**
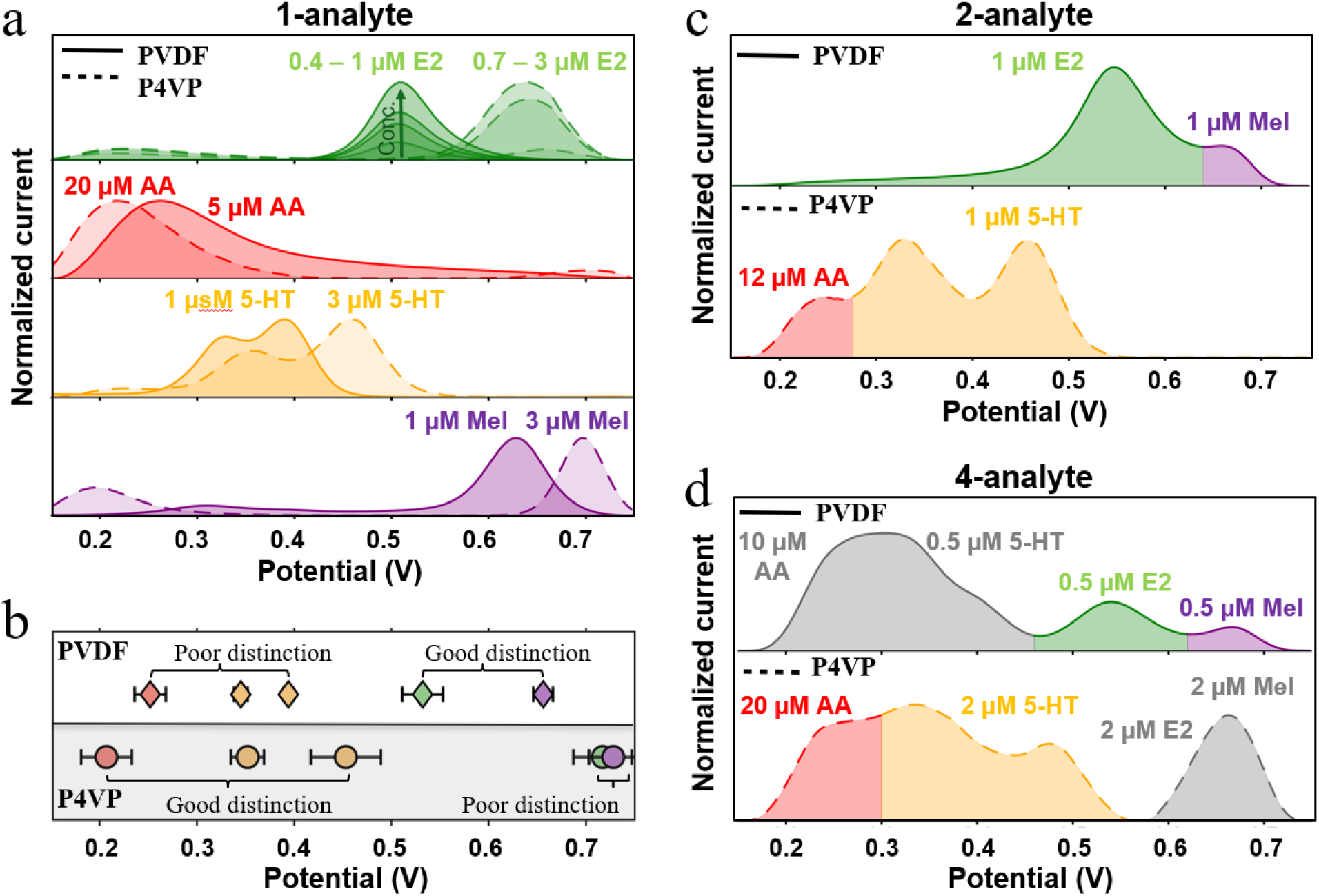
— Selective Detection Principle. **(a)** Normalized SWV data of PVDF-CF and P4VP-CF for E2, AA, 5-HT, and Mel as single analytes (three different concentrations for E2). **(b)** Oxidation peak potentials, showing better distinction of E2 (green) and Mel (purple) with PVDF-CF, while P4VP-CF provides better separation of AA (red) and 5-HT (yellow). Error bars represent three separately fabricated sensors. **(c)** Pairwise and **(d)** four-analyte mixture voltammograms.

Similar trends are observed for 5-HT and Mel: oxidation peaks appear at lower potentials with PVDF-CF, while P4VP-CF shifts them to higher values. Notably, 5-HT exhibits two oxidation peaks, which may originate from first oxidation to the carbocation, followed by oxidation to quinone imine, as previously reported.^14^ Interestingly, AA exhibits reversed behavior: P4VP-CF shifts the peak to lower potentials, while PVDF-CF shifts it to higher values. Note that while all measurements were conducted in PBS, the behaviors for estradiol detection in artificial saliva and under varying salt levels appear similar as reported in our previous work.^28^

These polymer-induced potential shifts form the basis for quantitative electrochemical analysis and enable ML models to distinguish and deconvolute overlapping signals in mixtures. The relative separation of analyte peak positions differs markedly between the two polymers, as summarized in **Fig. 2b**. On PVDF-CF, AA and 5-HT peaks appear closer together, while E2 and Mel are well separated. Conversely, P4VP-CF provides clear separation between AA and 5-HT, while E2 and Mel peaks overlap. This complementary behavior highlights the orthogonal selectivity achieved through polymer modification.

The importance of these findings becomes more evident in analyte mixture measurements. In 2-analyte mixtures, PVDF-CFs resolve E2 and Mel as two separate peaks, consistent with single-analyte measurements (**Fig. 2c**, raw data **Fig. S3e**). Similarly, P4VP-CFs separate AA and 5-HT signals. However, in 4-analyte mixtures, PVDF-CFs cannot distinguish AA and 5-HT, while P4VP-modified electrodes show overlapping signals for E2 and Mel (**Fig. 2d**, raw data **Fig. S3f**). While basic regression approaches may suffice for the examples shown in Fig. 2, deconvolution becomes increasingly challenging when analytes are present at vastly different concentration levels, as commonly occurs in physiological samples (e.g., when vitamin C is supplemented). This motivates the use of ML models, which can learn and exploit polymer-dependent peak shifts and signal shapes to enable robust multiplexed biomarker detection.

### Performance and Model Comparison

Fig. 3 summarizes the predictive performance of the proposed ML framework across analytes, error metrics, and model architectures using the derivative preprocessing on the combined data set. **Fig. 3a–d** show predicted versus experimentally measured concentrations obtained using the best-performing MLP model on a representative train-test split (linear axis shown in **Fig. S4a–d**). To reduce false non-zero predictions, classifier-gated masking was applied in zero-analyte cases (**Fig. S4e**). This masking resulted in a modest performance improvement (i.e., test *R*^2^ increased from 0.969 to 0.972 for Mel) and was therefore applied only for the analysis shown in Fig. 3. For all four analytes, predictions closely follow the identity line, yielding coefficients of determination (*R*^2^) exceeding 0.91. The slightly lower *R*^2^ observed for AA likely reflects the reduced sensitivity of the sensor towards this biomarker. For this representative split, the model achieves an overall *R*^2^ of 0.947 averaged across biomarkers. A comprehensive multi-seed comparison across all six regression models and analytes, reporting mean ± standard deviation across five random seeds, is provided in **Table S4** and a summary is shown in **Table 1**. RF and XGB showed larger train–validation *R*^2^ gaps (i.e., higher training *R*^2^ with reduced validation/test *R*^2^), consistent with overfitting, whereas the MLP exhibited a smaller generalization gap while achieving the best overall test metrics.

**Figure 3:**
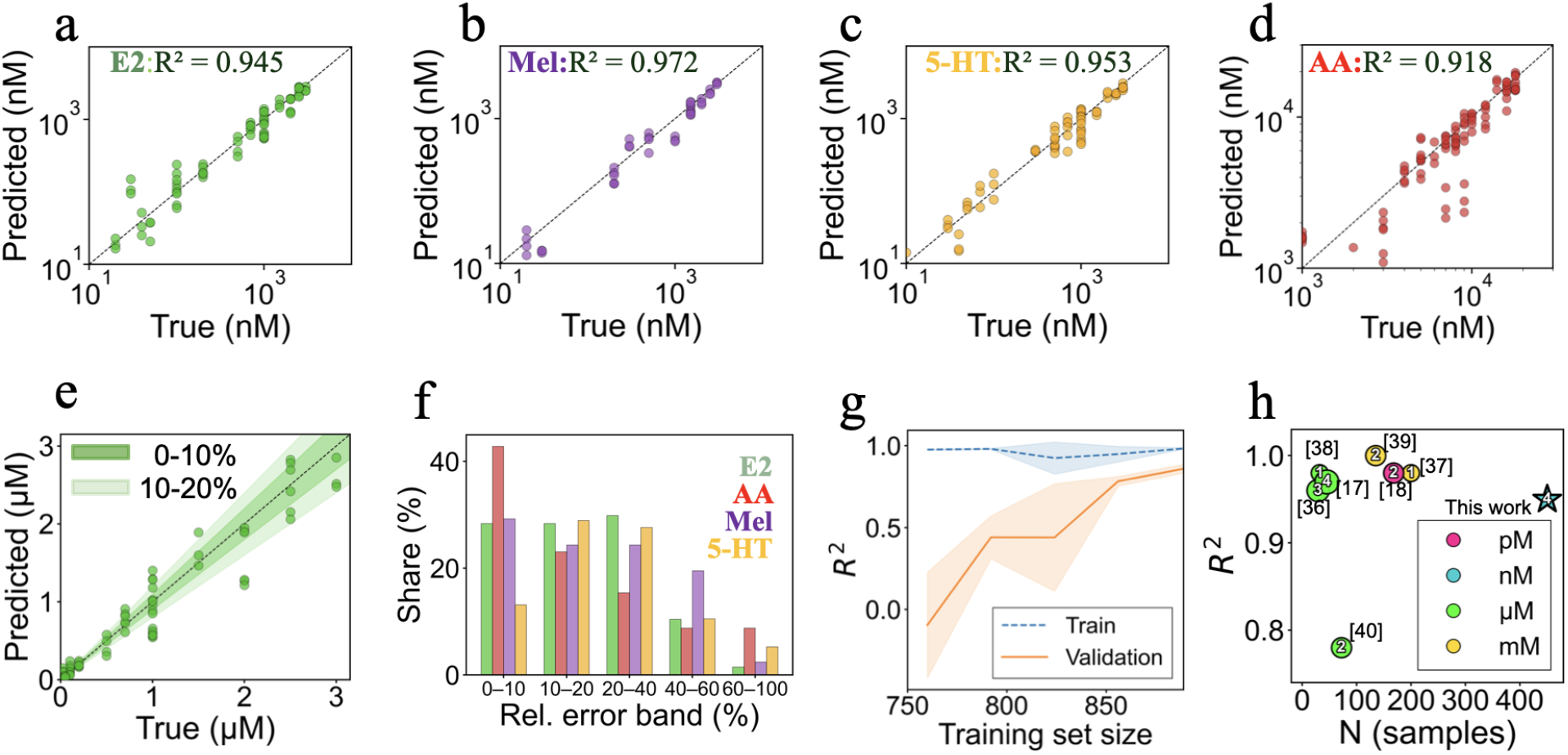
— Key ML Performance. Predicted versus true concentrations (log-log) and *R*^2^ values for a representative MLP model (seed = 42) for **(a)** E2, **(b)** Mel, **(c)** 5-HT, and **(d)** AA. For this representative split, a zero-aware two-stage architecture (classifier→ regressor) was used and regression outputs were masked to zero when the classifier predicted analyte absence, to suppress false non-zero predictions. **(e)** Relative MAE bands (0-10% and 10-20%) for E2 predictions. **(f)** Distribution of relative prediction errors across analytes. **(g)** Learning curve of the MLP model showing training and validation *R*^2^ as a function of training set size. **(h)** Comparison of this work with previously reported electrochemical ML sensing studies; note that reported *R*^2^ metrics may refer to calibration, cross-validation, or held-out test sets depending on the study (see Table S5)

**Table 1:** Overall model performance. Test metrics averaged across all four analytes and five random seeds using derivative preprocessing on the combined dataset. Best split: MLP seed 42 with classifier-gated masking.

| Model | $R^2$ | MAE (nM) | RMSE (nM) |
| --- | --- | --- | --- |
| Ridge | 0.611 | 1020 | 1441 |
| RF | 0.828 | 492 | 799 |
| KNN | 0.828 | 453 | 769 |
| XGBoost | 0.848 | 475 | 751 |
| SVR | 0.892 | 414 | 696 |
| **MLP** | **0.908** | **409** | **644** |
| MLP (best split) | 0.947 | 312 | 542 |

Robust prediction accuracy is maintained down to tens to hundreds of nanomolar for estradiol and serotonin (**Fig. S4f-h**), demonstrating stable performance across a wide dynamic range. While further sensitivity gains would be advantageous for non-invasive measurements in sweat and saliva (sometimes required sensitivity is in the pM range), the current results already indicate strong predictive capability. Importantly, these data were acquired using multiple sensors operated over days to weeks without calibration: As shown in **Fig. S5**, noticeable inter-sensor variations are observed in absolute peak currents, likely arising from differing electrochemical surface areas. In contrast, oxidation potentials show only minimal variation (i.e., 0.532 ± 0.026 V for PVDF-CF and 0.679 ± 0.002 V for P4VP-CF), highlighting that the polymer-induced shift is highly reproducible. Over an operation period of 37 days, we observe a reduced E2 peak current (from 2.45 to 0.84 nA) and an increase in oxidation potential (i.e., from 0.52 to 0.60 V (**Fig. S6**). The near-perfect overlap between electrodes on the same substrate from the same fabrication batch (**Fig. S7**) supports the use of a single electrode per sensor substrate for ML training. The capability of the ML model to accurately predict analyte concentrations despite inter-sensor variabilities and temporal drift further underscores the robustness of both the sensor platform and the ML framework. This is further supported by a noise robustness analysis showing that the model maintains *R*^2^ *>* 0.79 for all analytes up to 2% additional noise when SG smoothing is applied (**Fig. S8**).

In addition to *R*^2^, prediction accuracy was evaluated using the mean absolute error (MAE), shown exemplarily for E2 in **Fig. 3e** for the 0–10% and 10–20% error bands. MAE provides a complementary, scale-aware measure of absolute prediction error and enables intuitive interpretation of quantitative accuracy. Approximately 30% of predictions fall within the 0–10% band, with an additional 30% within the 10–20% range. The larger errors are likely attributed to the complexity of the system (i.e., testing analytes with similar oxidation potentials and varying concentration ranges), as well as the inter-sensor variability and stability over time. The combination of these factors reduces the accuracy of our sensor but represents a more realistic protocol. The relative error distributions across all biomarkers are summarized in **Fig. 3f**, with corresponding MAE values and standard deviations reported in **Fig. S4i**,**j**. For this representative split, relative errors exceeding 100% were observed only for E2 (4/71 non-zero samples). For AA, 5-HT, and Mel, the maximum relative error reached exactly 100%, arising from cases where negative regression outputs were clipped to zero. Although individual prediction errors can reach several tens of percent, the models consistently preserve concentration-dependent trends across all biomarkers. This is an outcome that is hardly achievable using conventional peak-based or univariate analysis for datasets and biomarkers with such strongly overlapping voltammetric signals. Importantly, physiological variations in these biomarkers frequently span one to two orders of magnitude (e.g., day–night melatonin levels vary by up to 100×), indicating that the achieved accuracy is sufficient to resolve biologically meaningful changes.

Comparing our work to the state of the art is inherently challenging, as previous studies have applied ML to electrochemical sensors under highly variable conditions, including differences in number of biomarkers, sample size, sensor type, and measurement environment. To enable a meaningful comparison, we focus on four key parameters: sample size, number of biomarkers, *R*^2^, and targeted concentration range (**Fig. 3h**, details in **Table S5**). Our work stands out by introducing a dataset that is both substantially larger (450 samples, nearly an order of magnitude greater than most prior ML studies) and more diverse, encompassing a wide concentration range (down to nM) and complex mixtures (up to four biomarkers), while achieving comparable *R*^2^ values. This was accomplished despite deliberately selecting biomarkers that oxidize within a narrow potential window (0.2–0.8 V), which increases the challenge of signal deconvolution. It is important to note, however, that some prior studies were conducted in real human fluids, which introduce additional complexity not addressed in our current work.

Overall, we attribute the strong performance of our system to the use of polymer-diverse electrode modifications, which enhance peak separation and enable more effective signal deconvolution, as shown in Fig. 2. Moreover, rigorous validation, including cross-validation and evaluation across five random seeds, ensures a robust assessment of model generalizability. Learning curve analysis (**Fig. 3g**) indicates that the MLP model’s test *R*^2^ has not yet plateaued with respect to dataset size, suggesting that additional training data could further improve predictive accuracy. Potential systematic prediction bias was further assessed using residuals as a function of true concentration for melatonin as a representative example (**Fig. S4l**). In summary, our work combines polymer-modified sensors with a large, mixture-rich dataset and multi-seed validation, enabling a robust evaluation of multiplexed electrochemical ML performance and providing a foundation for studies in more complex, physiologically relevant fluids.

### Effect of Polymer: Feature Analysis

To explore the importance of polymer modification, we trained the optimized models separately on PVDF-CF and P4VP-CF datasets and compared their performance (**Fig. 4** and **Table S6**). All models were tuned using identical hyperparameter search spaces and evaluated without applying a classification mask, ensuring a fair and consistent comparison. Note that the goal of Fig. 4 is to illustrate polymer effects rather than focusing on absolute performance, as evaluating polymers separately halves the dataset, and data splits reduce absolute *R*^2^.

**Figure 4:**
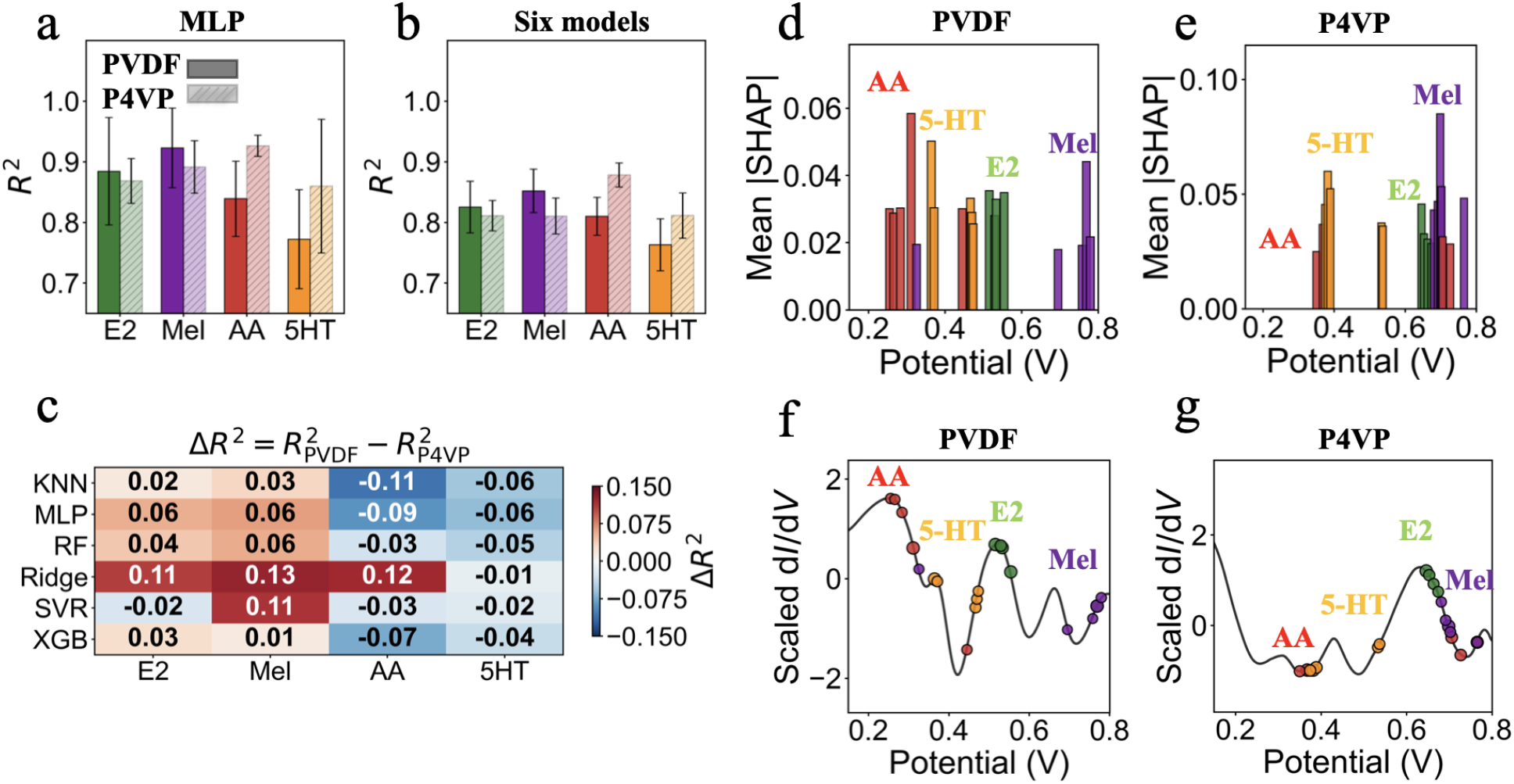
— Feature Importance and Interpretability. **(a)** *R*^2^ values for derivative preprocessing for the MLP model only and **(b)** averaged across all six models (*R*^2^ *>* 0.6). **(c)** Heatmap of 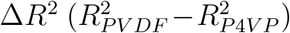 across all preprocessing transformations (*R*^2^ > 0.6). Red shades indicate better performance for PVDF, while blue shades indicate better performance for P4VP **(d)** Top 5 SHAP values of all analytes for the PVDF-CF and **(e)** P4VP-CF data using derivative transformed signals. **(f)** Top 5 SHAP values mapped onto the average derivative signal of PVDF-CF and **(g)** P4VP-CF data using the MLP model.

The polymer-specific *R*^2^ values obtained using the MLP model are shown in **Fig. 4a**. PVDF-CF outperforms P4VP-CF for predicting E2 and Mel, whereas P4VP yields superior performance for AA and 5-HT, in excellent agreement with the electrochemical observations in Fig. 2. This polymer-dependent selectivity persists across all ML models (**Fig. 4b**). The heatmap (**Fig. 4c**) summarizes the performance difference 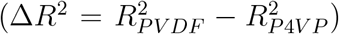 and reveals the same consistent trend: PVDF-CF is advantageous for the detection of E2 and Mel (red), while P4VP-CF outperforms PVDF-CF for AA and 5-HT (blue), with only a few exceptions. Importantly, **Table S6** reports the complete set of *R*^2^ values without applying any threshold, ensuring full transparency of model performance. Additionally, the heatmap represents data averaged across the three preprocessing steps, including derivative (used in Figs. 3 and 4), min-centering, and polynomial. Min-centering and polynomial perform slightly inferiorly to the derivative method, which is why the derivative was chosen throughout most of our study; however, as shown in the heatmap and supporting table, the polymer-dependent selectivity trends hold true across all the preprocessing methods, demonstrating that the preprocessing does not alter the underlying electrochemical information. Note that in Fig. 4a-c, only polymer–biomarker combinations exceeding a threshold of *R*^2^ *>* 0.6 are displayed to emphasize robust and practically relevant predictions. Overall, our results highlight that polymer modification is a key factor in enhancing sensor performance, with the close agreement between electrochemical observations and ML results confirming that polymer-induced peak separation directly improves predictive accuracy.

To further investigate polymer-specific selectivity, we applied SHAP to the MLP model to assess the feature importance of each polymer dataset. SHAP quantifies the contribution of individual features, ^33^ in our case current–potential points, enabling direct mapping between learned model features and regions of the electrochemical signal. **Fig. 4d**,**e** show the top five voltage features ranked by mean absolute SHAP value for all analytes using the P4VP-CF and PVDF-CF datasets with derivative-transformed traces (additional SHAP values in **Fig. S9**). The MLP model relies on distinct voltage regions for different analytes, and changing the polymer coating shifts which regions are deemed informative; specifically, E2 and Mel shift to higher potentials on P4VP-CF and can only be distinguished on PVDF-CF, whereas AA and 5-HT shift to lower and higher potentials, respectively, on P4VP-CF, enabling their distinction, again consistent with Fig. 2.

Mapping the most influential features onto average signal traces (**Fig. 4f**,**g**) shows that key ML features correspond to extrema, zero crossings, and high-slope regions, i.e., regions of analyte oxidation. Although these results are based on derivative-transformed data and absolute potentials may appear shifted relative to Fig. 2, the same trends hold for polynomial-transformed signals (**Fig. S10**), confirming that the model relies on distinct potential regions for different analytes depending on the polymer coating.

Overall, the feature analysis and SHAP results underscore the critical role of polymer modifications in multiplexed sensing, confirming that orthogonal sensor arrays are essential for effective ML deconvolution of overlapping voltammetric signals; something a single sensor type cannot achieve. Crucially, our findings provide a mechanistic explanation for how polymer-induced selectivity enhances multiplexed sensing, rather than applying ML as a black box—a significant advance, as many studies do not clarify why their models succeed. ^34^ This work demonstrates how interpretability tools can connect electrochemical intuition with data-driven ML models. It also highlights the potential of a two-electrode array for four analytes and provides a roadmap for scaling to more complex datasets by incorporating additional polymer coatings that induce distinct peak shifts.

#### Limitations and Outlook

This study demonstrates that a two-electrode sensor array can effectively distinguish four analytes. However, several limitations must be addressed for translation to wearable applications. Estradiol and melatonin sensitivity needs to be enhanced to match physiological levels. Other molecules such as dopamine, epinephrine, norepinephrine, and most critically uric acid, feature oxidation potentials that overlap with the current molecules **(Fig. S11)**. Further evaluation under varying conditions (e.g., ionic strength, pH, artificial and real saliva) and their impact on the ML model is needed to improve robustness. Finally, our model is a closed-set regression trained on four analytes, and cannot detect unknown species in its current form, an inherent property of all closed-set ML sensor systems. We believe that, despite being demonstrated in a simplified system, the current work provides a viable strategy to address stringent selectivity limitations of electrochemical sensors and can enhance the robustness and selectivity of next-generation electrochemical sensors.

## Conclusion

This study presents an electrochemical sensor array capable of selectively detecting 5-HT, AA, E2, and Mel in mixture conditions. The array consists of carbon flower electrodes modified with PVDF or P4VP polymers and deposited onto stretchable substrates via a scalable process, making it well-suited for skin-based applications and readily adaptable to larger arrays. Polymer modifications shift analyte oxidation potentials, producing distinct, resolvable electrochemical signatures and generating high-quality datasets optimized for ML analysis. We evaluated 450 square wave voltammograms from single- and multi-analyte mixtures at clinically relevant concentrations. ML-based signal processing achieved an overall *R*^2^ of 0.947, with most MAE values within the 30% error band, sufficient to resolve biologically meaningful changes. Among six tested ML algorithms, the MLP model performed best, likely due to its ability to capture complex non-linear relationships in overlapping voltammetric signatures, outperforming linear models. Feature importance analysis revealed that PVDF-CFs are critical for distinguishing E2 and Mel, whereas P4VP-CFs enable discrimination between AA and 5-HT. SHAP analysis further confirmed that the models rely on oxidation-relevant regions, validating the mechanistic role of polymer-induced selectivity. In summary, this work demonstrates that polymer-modified carbon electrodes combined with interpretable ML can robustly discriminate and quantify multiple non-invasive biomarkers in complex mixtures. The approach offers a clear pathway for expansion to larger electrode arrays and broader analyte panels.

## Supporting Information

Preprocessing and features; SHAP sensitivity analysis; raw voltammograms; prediction accuracy and error analysis; inter-sensor variability; temporal drift; linearity and inter-electrode variability; noise robustness analysis with Savitzky–Golay filtering; SHAP density plot; SHAP feature importance; dataset; ML parameters; concentration ranges; model-dependent performance; literature comparison; preprocessing/polymer-dependent performance

## Acknowledgments

This project was supported by the Stanford Wearable Electronics Initiative (eWEAR) seed grant, the Tianqiao and Chrissy Chen Ideation and Prototyping Lab, the Swiss National Science Foundation (SNSF, I.C.W.: P500PT-214498), and the Turing Grant Scheme (A.L.M.D.). Part of this work was performed at the Stanford Nano Shared Facilities (SNSF), supported by the National Science Foundation under award ECCS-2026822. Z.B. is a Chan Zuckerberg Biohub San Francisco investigator and an Arc Institute innovation investigator. We thank L. Mondonico, A. Mahmud, K-J Hsu, and K. Parkatzidis for their scientific advice.

## Data and Code Availability

All electrochemical raw data, processed datasets, and analysis notebooks used in this study are available at the following: https://github.com/esemsc-ald24/ML-Biomarker-Sensing.gitGitHub repository.^35^ Further data are available from the corresponding author upon reasonable request.

## TOC Graphic

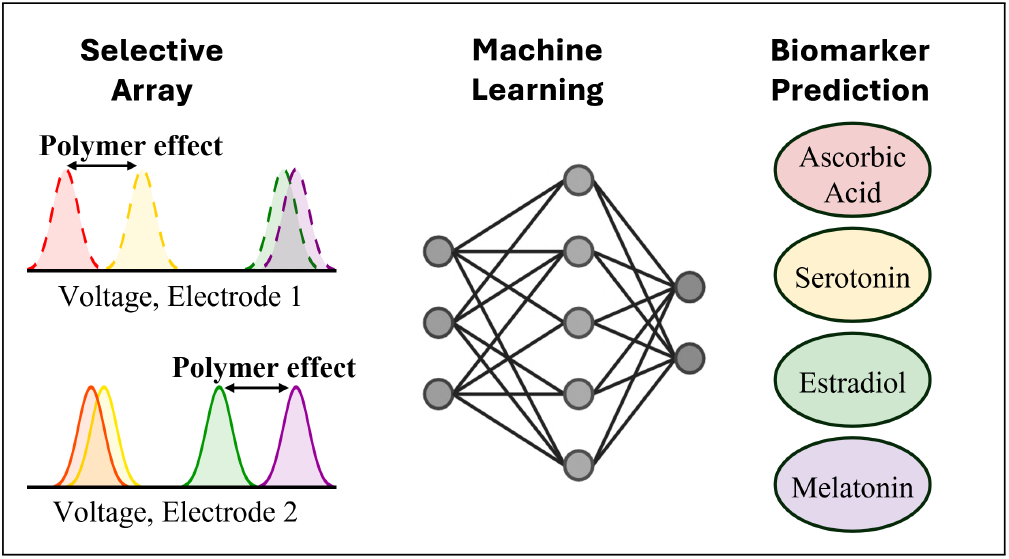

