## Supplementary Information for "Machine Learning-Driven Multiplexed Biomarker Detection with Polymer-Enhanced Electrochemical Sensors"

| | **Contents** |
|---|---|
| **Figure S1** | Preprocessing and Features |
| **Figure S2** | SHAP Sensitivity Analysis |
| **Figure S3** | Raw Voltammograms |
| **Figure S4** | Prediction Accuracy and Error Analysis |
| **Figure S5** | Inter-Sensor Variability |
| **Figure S6** | Temporal Drift of PVDF-CF |
| **Figure S7** | Linearity and Inter-Electrode Variability |
| **Figure S8** | Noise Robustness Analysis with Savitzky–Golay Filtering |
| **Figure S9** | SHAP Density Plot |
| **Figure S10** | SHAP Feature Importance |
| **Figure S11** | Additional Biomarkers |
| **Table S1** | Dataset |
| **Table S2** | ML Parameters |
| **Table S3** | Concentration Ranges |
| **Table S4** | Model-Dependent Performance |
| **Table S5** | Literature Comparison |
| **Table S6** | Preprocessing/Polymer-Dependent Performance |

### Supplementary Figures

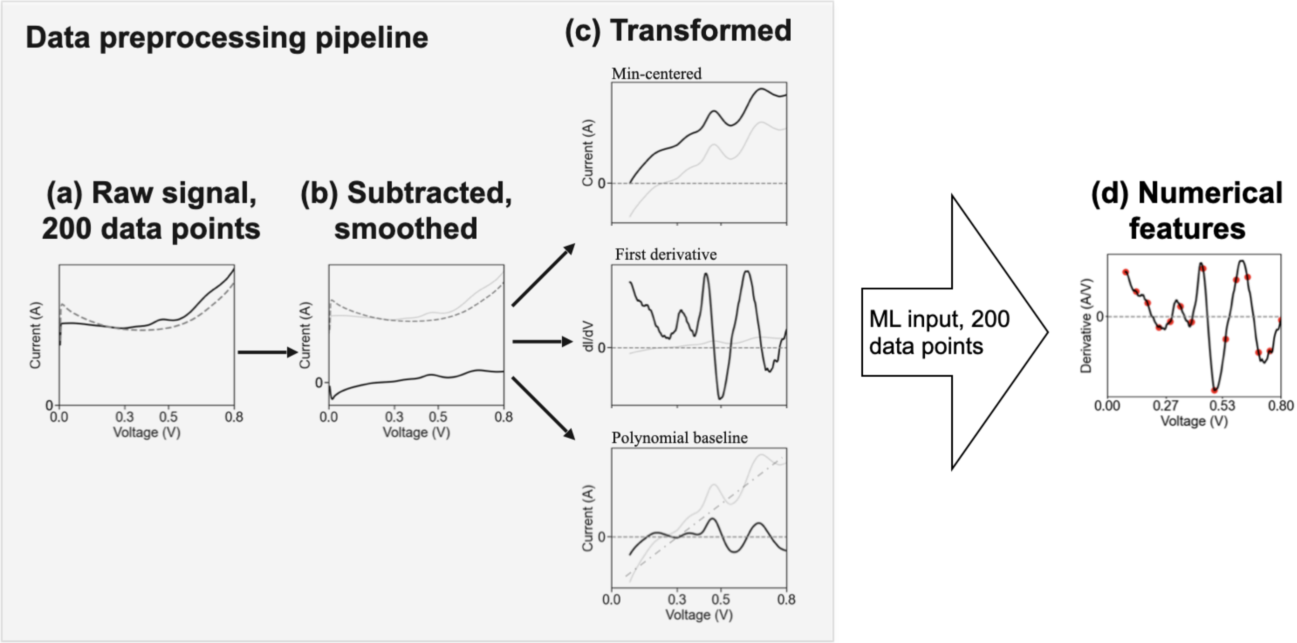


**Figure S1.** | **Preprocessing and Features.** **(a)** Raw P4VP-CF signal of a mixture containing 500 nM E2, 9 µM AA, and 2000 nM 5-HT, shown together with the corresponding blank PBS signal (dashed). **(b)** Blank PBS subtraction followed by smoothing. **(c)** Signal transformations, including min-centering, polynomial baseline subtraction, and first-derivative transformation ($dI/dV$). **(d)** Numerical feature representation used as ML input, consisting of the full transformed current trace with 200 data points per scan (only 20 points shown for clarity). For visualization purposes, signals below 0.1 V are omitted in panels (c) and (d); the full potential range (0.0–0.8 V) was used for all analyses.

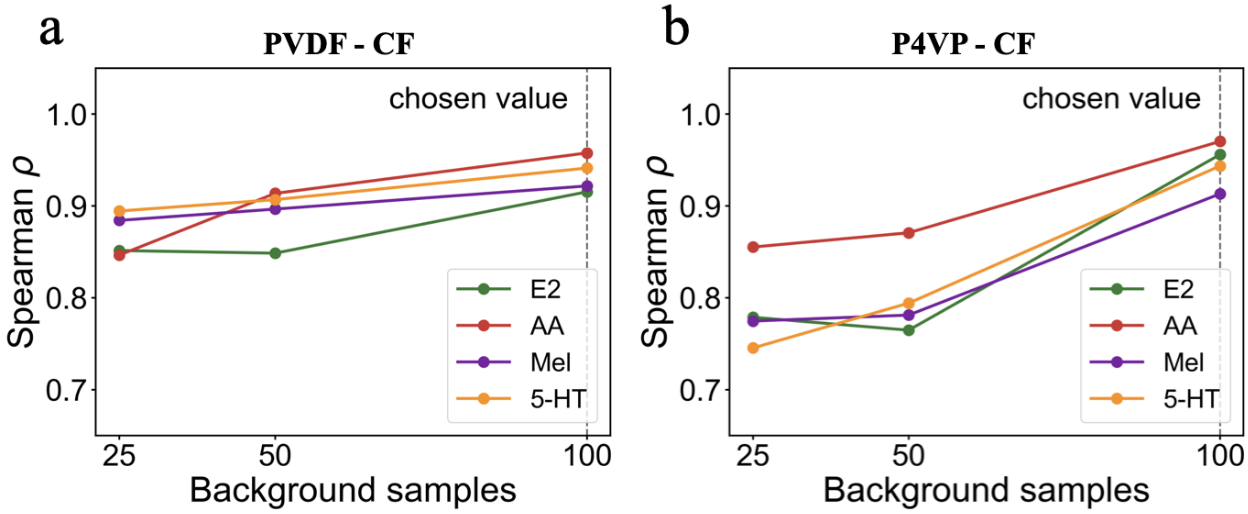


**Figure S2.** | **SHAP Sensitivity Analysis.** SHAP ranking stability for PVDF-CF **(a)** and P4VP-CF **(b)**, assessed via Spearman correlation between rankings obtained with varying background sample sizes (n = 25, 50, 100). Dashed line indicates the chosen value of n = 100; all analytes converge above $\rho = 0.9$ at this threshold, confirming stable feature rankings.

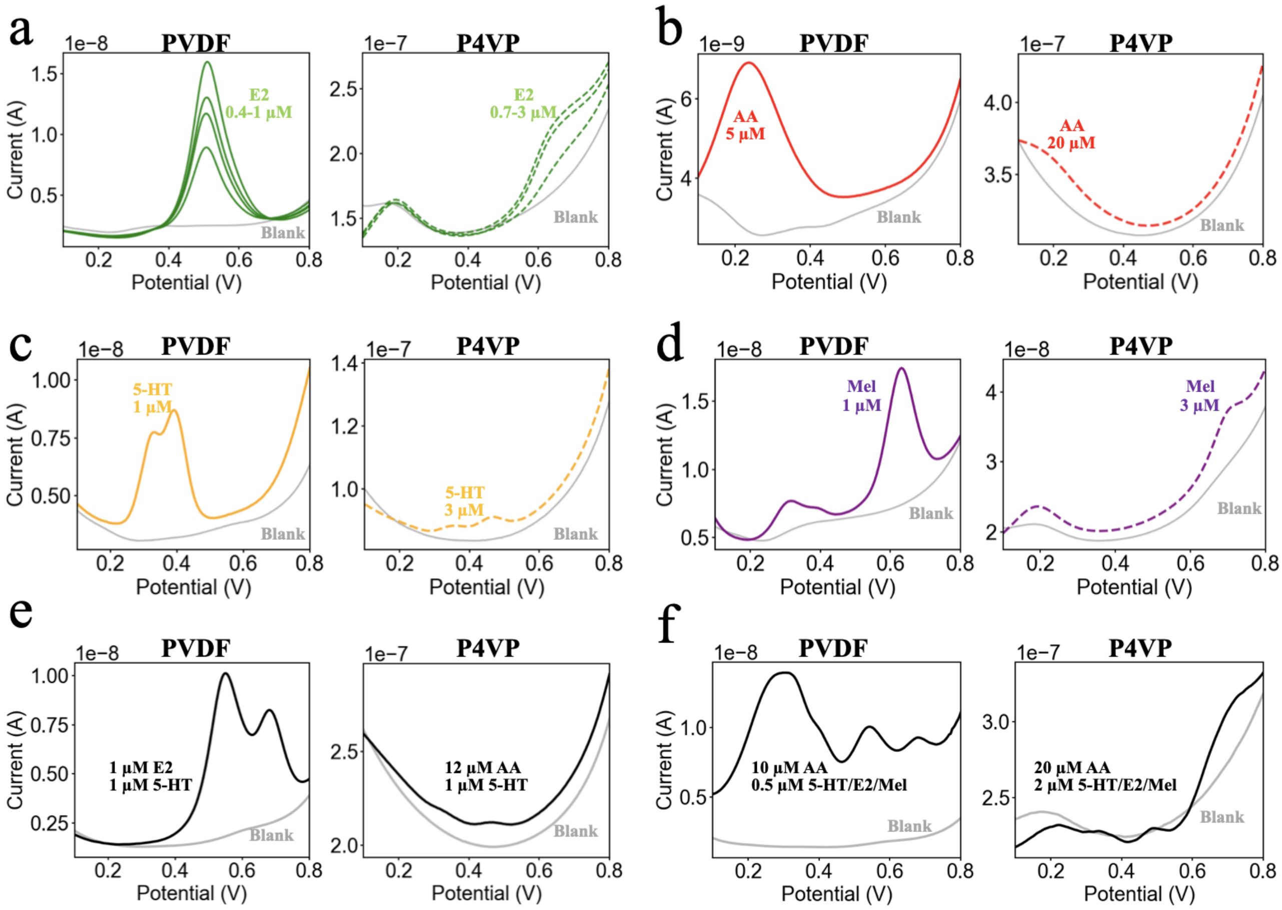


**Figure S3. | Raw Voltammograms. (a)** Raw E2 single-analyte voltammograms (including blank PBS) using PVDF-CF and P4VP-CF. **(b-d)** Analogous data for AA, 5-HT, and Mel, respectively. **(e)** Raw two-analyte voltammograms (including blank PBS) using PVDF-CF and P4VP-CF. **(f)** Raw four-analyte voltammograms (including blank PBS) using PVDF-CF and P4VP-CF.

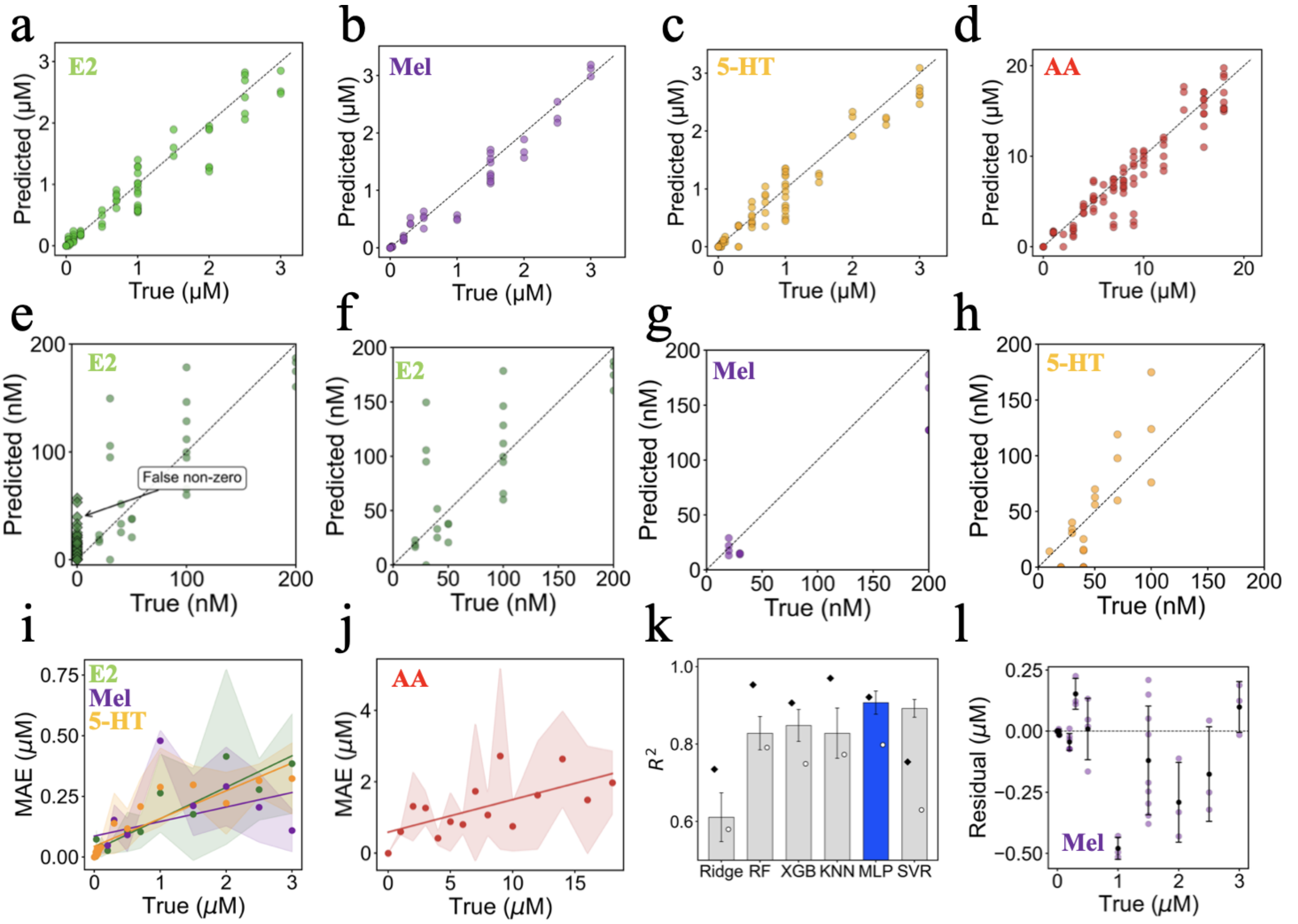


**Figure S4.** | **Prediction Accuracy and Error Analysis.** **(a–d)** Predicted versus true concentrations (linear). **(e)** False non-zero predictions for E2, illustrating cases where the regression model predicts finite concentrations in the absence of the analyte. **(f–h)** Predicted versus true concentrations for E2, Mel, and 5-HT in the low-concentration regime. **(i)** MAE with standard deviation across seeds for E2, Mel, and 5-HT. **(j)** MAE with standard deviation across seeds for AA. **(k)** Model comparison across all six regression models; bars show test $R^2$, black diamonds indicate training $R^2$, and white circles indicate validation $R^2$. **(l)** Residuals as a function of true concentration for melatonin, used to assess systematic prediction bias. All panels show results for the MLP model using a representative split (seed = 42).

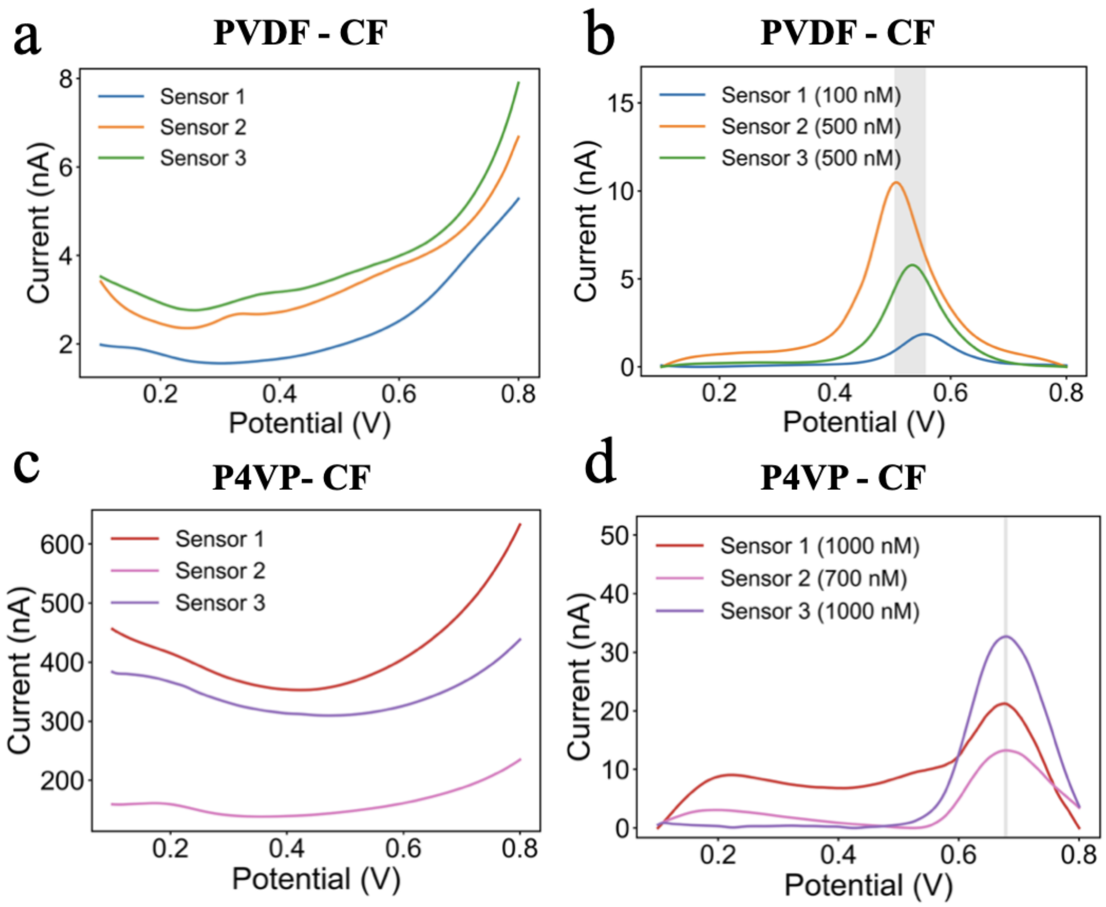


**Figure S5. | Inter-Sensor Variability. (a)** PBS blank SWV voltammograms of three PVDF-CF sensors. **(b)** Background-subtracted E2 detection with PVDF-CF (note the different concentrations). **(c)** PBS blank SWV voltammograms of three P4VP-CF sensors. **(d)** Background-subtracted E2 detection with P4VP-CF (note the different concentrations). The grey shaded region indicates the range of observed E2 oxidation potentials.

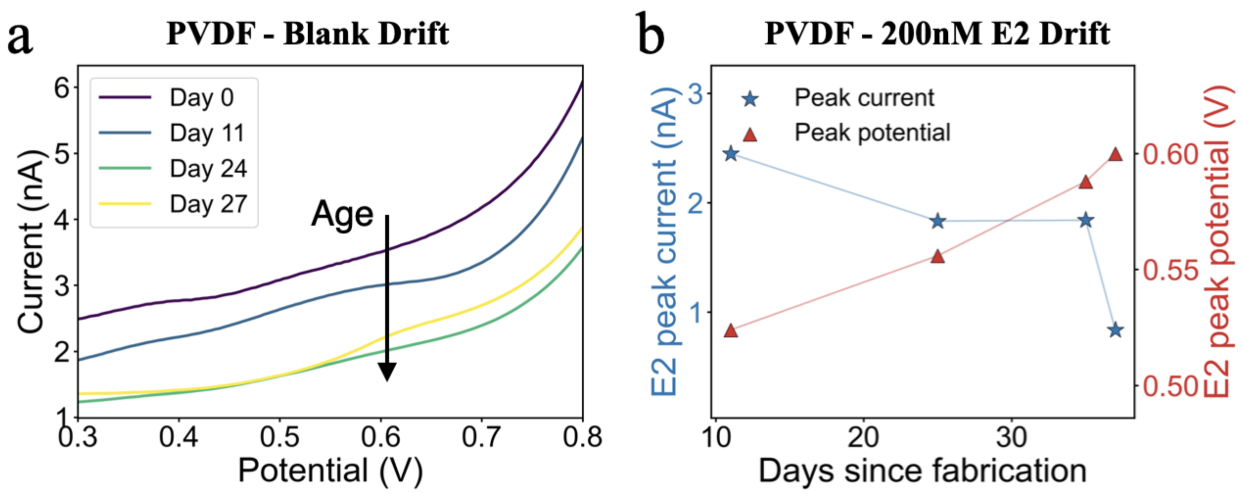


**Figure S6. | Temporal Drift of PVDF-CF. (a)** Blank PBS voltammograms recorded with the same electrode (MC325) on days 0, 11, 24, and 27. **(b)** Estradiol peak current (stars, left ordinate) and potential (triangle, right ordinate) after blank subtraction as a function of sensor age .

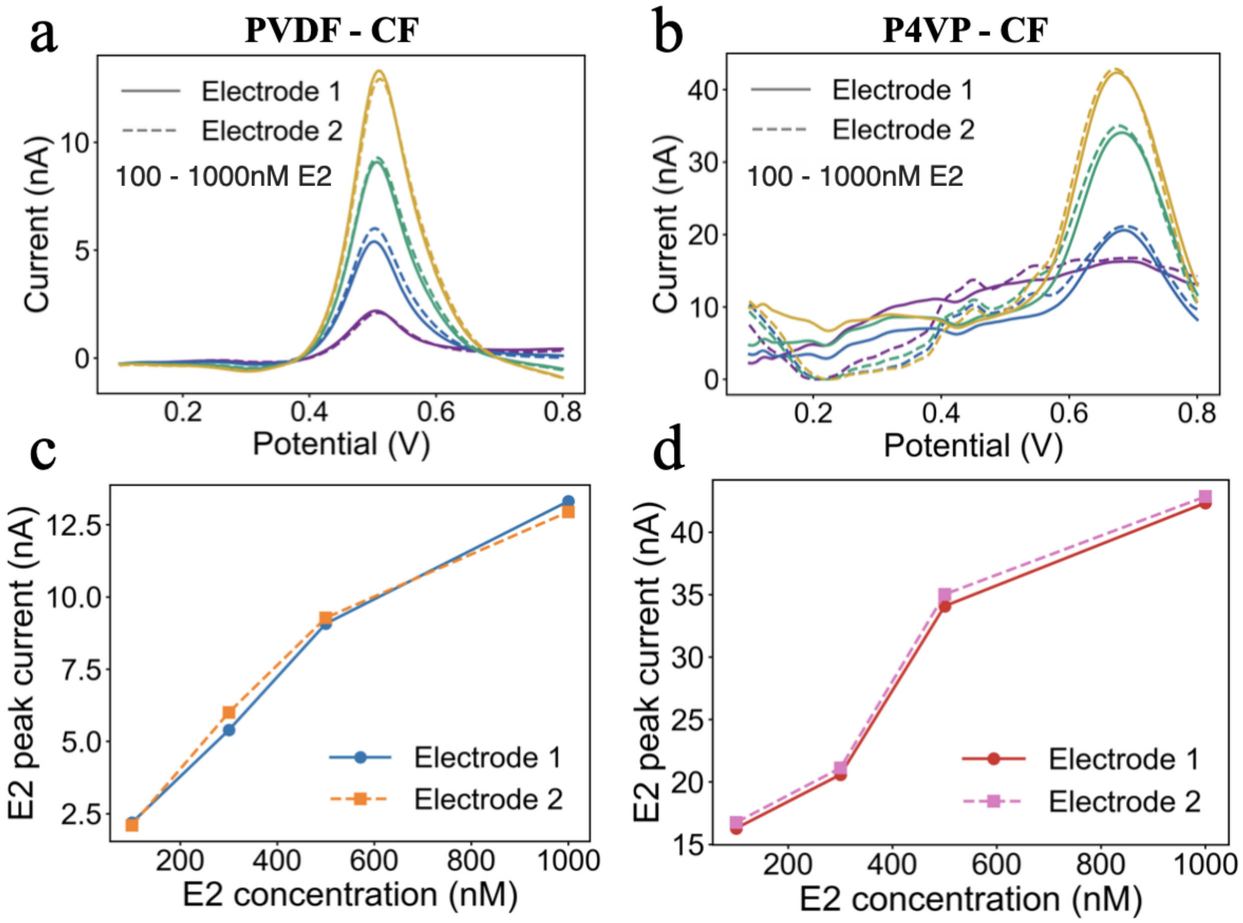


**Figure S7.** | **Linearity and Inter-Electrode Variability.** **(a)** Background-subtracted E2 voltammograms on PVDF-CF at 100–1000 nM, comparing two parallel electrodes (solid vs dashed). **(b)** Analogous on P4VP-CF. **(c,d)** Calibration curves showing E2 peak current vs. concentration for both electrodes on PVDF-CF (c) and P4VP-CF (d).

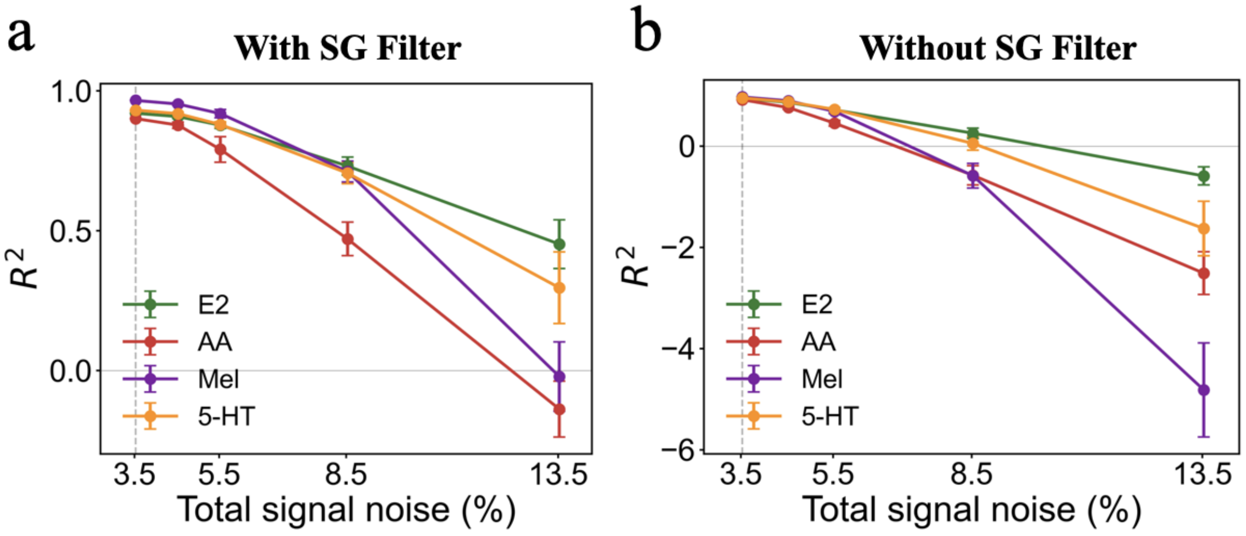


**Figure S8.** | **Noise Robustness Analysis with Savitzky–Golay Filtering. (a)** $R^2$ vs. added Gaussian noise (0–10% of signal standard deviation) with SG smoothing. The baseline (0%) already includes 3.5% instrument noise (SNR = 276). (b) Analogous without SG smoothing. Error bars denote standard deviation over 10 random noise seeds.

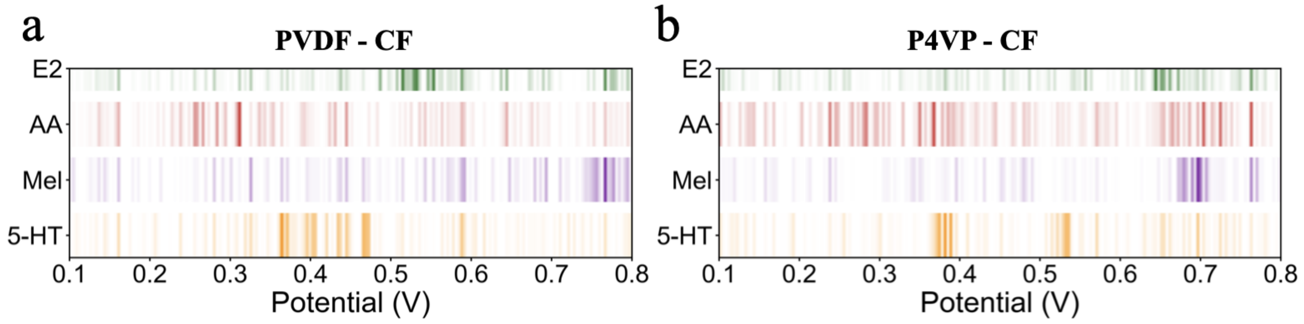


**Figure S9.** | **SHAP Density Plot.** Normalized SHAP importance mapped across the voltage range for each analyte on the PVDF **(a)** and P4VP **(b)** sensors. Color intensity indicates relative feature importance for predicting the respective analyte concentration.

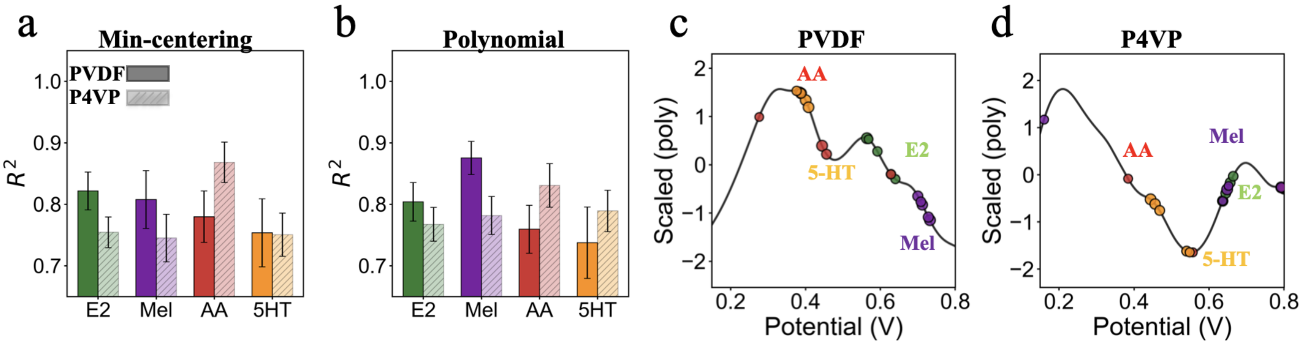


**Figure S10.** | **Feature Importance. (a)** $R^2$ values for min-centering preprocessing averaged across all models. **(b)** Analogous data for polynomial baseline preprocessing. **(c)** Top five SHAP values mapped back onto the average polynomial-subtracted signal of PVDF-CF, **(d)** and P4VP-CF data sets using the best MLP model for all analytes.

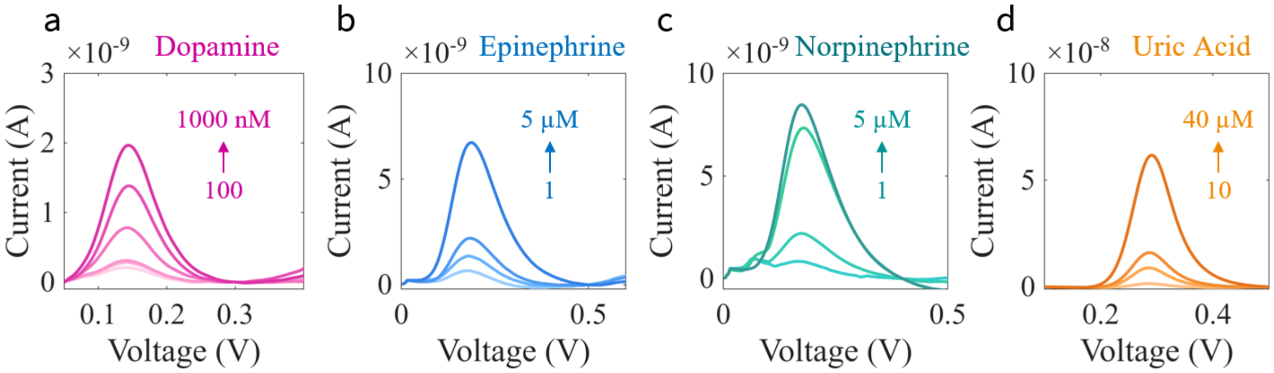


**Figure S11.** | **Additional Biomarkers.** SWV detection of dopamine **(a)**, epinephrine **(b)**, norepinephrine **(c)**, and uric acid **(d)** with a PVDF-CF electrode.

### Supplementary Tables

**Table S1. Dataset.** Overview of the complete dataset, indicating the polymer, sensor ID, biomarkers, and number of unique measurements. The dataset includes at least 10 samples for each individual analyte and 15 samples for 2-, 3-, and 4-analyte mixtures (both for PVDF-CF and P4VP-CF), totalling 450 samples.

| Polymer | Sensor ID | Biomarker | # Measurements | Total |
|---|---|---|---|---|
| P4VP | MC302 | E2 | 6 | **6** |
| P4VP | MC312 | AA | 5 | **5** |
| P4VP | MC327 | E2 | 6 | |
| | | AA | 10 | |
| | | 5-HT | 6 | |
| | | Mel | 6 | |
| | | E2/AA | 15 | |
| | | E2/5-HT | 15 | |
| | | E2/Mel | 15 | |
| | | AA/5-HT | 15 | |
| | | AA/Mel | 15 | **103** |
| P4VP | MC329 | E2 | 4 | |
| | | 5-HT | 4 | |
| | | Mel | 4 | |
| | | 5-HT/Mel | 15 | |
| | | E2/AA/5-HT | 15 | |
| | | E2/AA/Mel | 15 | |
| | | AA/5-HT/Mel | 15 | **72** |
| P4VP | MC330 | E2/5-HT/Mel | 15 | |
| | | E2/AA/Mel | 4 | |
| | | E2/AA/5-HT/Mel | 15 | **34** |
| PVDF | MC313 | E2 | 5 | |
| | | E2/AA | 5 | |
| | | E2/AA/5-HT/Mel | 1 | **11** |
| PVDF | MC320 | E2 | 13 | |
| | | AA | 10 | |
| | | 5-HT | 13 | |
| | | Mel | 13 | |
| | | E2/Mel | 1 | **50** |
| PVDF | MC325 | AA | 5 | |
| | | E2/AA | 15 | |
| | | E2/5-HT | 15 | |
| | | E2/Mel | 14 | |
| | | AA/5-HT | 15 | |
| | | AA/Mel | 15 | |
| | | 5-HT/Mel | 15 | |
| | | E2/AA/5-HT | 15 | |
| | | E2/AA/Mel | 15 | |
| | | E2/5-HT/Mel | 15 | |
| | | AA/5-HT/Mel | 15 | |
| | | E2/AA/5-HT/Mel | 15 | **169** |
| | | | | **Total = 450** |

**Table S2. ML Parameters.** Summary of parameters and tools used. The most frequently selected values across seeds are indicated in bold.

| **Category** | **Parameter (Unit)** | **Value / Setting** |
|---|---|---|
| **Hyperparameter Search Grids** | Ridge $\alpha$ | {0.1, 1.0, **10.0**, 100.0} |
| | RF n_estimators | {100, **200**} |
| | RF max_depth | {5, **10**} |
| | RF min_samples_leaf | {**1**, 3} |
| | MLP hidden_layer_sizes | {(64), (128), (128,64), (256,128), **(256,128,64)**} |
| | MLP learning_rate_init | {0.0003, **0.001**, 0.003} |
| | MLP $\alpha$ (L2 penalty) | {$1\times10^{-5}$, $\mathbf{1}\times10^{-4}$, $1\times10^{-3}$} |
| | KNN n_neighbors | {3, **5**, 7} |
| | KNN weights | {uniform, **distance**} |
| | XGBoost n_estimators | {100, **200**} |
| | XGBoost learning_rate | {0.01, **0.1**} |
| | XGBoost max_depth | {**3**, 5} |
| | SVR C | {0.1, 1.0, **10.0**} |
| | SVR $\epsilon$ | {**0.01**, 0.1} |
| | SVR kernel | {linear, **rbf**} |
| **Software** | Core ML Libraries | scikit-learn $\geq$ 1.2, XGBoost $\geq$ 1.6 |
| | Data Processing | NumPy $\geq$ 1.23, Pandas $\geq$ 1.5, SciPy $\geq$ 1.9 |
| | Visualization | Matplotlib $\geq$ 3.6, Seaborn $\geq$ 0.12 |
| | Model Interpretation | SHAP $\geq$ 0.41 |
| | Statistical Analysis | Statsmodels $\geq$ 0.13 |
| | Utilities | tqdm $\geq$ 4.64 |
| | Development Environment | JupyterLab $\geq$ 3.4, pytest $\geq$ 7.0, MkDocs + Material theme |

**Table S3. Concentration Ranges.** Comparison of physiological biomarker levels in sweat and saliva with the concentrations used for the sensor.

| Molecule | Sweat | Saliva | Sensor |
|---|---|---|---|
| Ascorbc acid | 0.01–50 µM [1] | 5-10 µM [2] | 1-20 µM |
| Serotonin | | 20–300 nM [3] | 10-3000 nM |
| Estradiol | 0-50 pM [4] | 0-50 pM [5] | 10-3000 nM |
| Melatonin | | 20-500 pM [6] | 10-3000 nM |

**Table S4. Model-Dependent Performance.** Performance evaluation for each biomarker (mean ± standard deviation) reporting **test** $R^2$, **MAE**, and **RMSE** (nM; note AA concentrations are in the µM range, reported here in nM for consistency). Overall model performance, averaged across biomarkers, is also reported and includes test, train, and validation $R^2$, as well as model training time. Models were evaluated with five random seeds using a held-out 10% test set, without classifier-gated masking. The MLP achieves the best overall performance.

**$R^2$ (test set)**

| Model | E2 | AA | Mel | 5-HT |
|---|---|---|---|---|
| Ridge | 0.671 ± 0.046 | 0.400 ± 0.152 | 0.705 ± 0.102 | 0.666 ± 0.070 |
| RF | 0.850 ± 0.064 | 0.845 ± 0.059 | 0.839 ± 0.078 | 0.779 ± 0.060 |
| XGBoost | 0.853 ± 0.040 | 0.867 ± 0.041 | 0.852 ± 0.077 | 0.820 ± 0.056 |
| KNN | 0.847 ± 0.112 | 0.865 ± 0.036 | 0.847 ± 0.050 | 0.752 ± 0.116 |
| **MLP** | **0.932 ± 0.027** | **0.888 ± 0.056** | **0.943 ± 0.023** | **0.867 ± 0.066** |
| SVR | 0.914 ± 0.030 | 0.874 ± 0.028 | 0.910 ± 0.033 | 0.870 ± 0.058 |

**MAE (nM)**

| Model | E2 | AA | Mel | 5-HT |
|---|---|---|---|---|
| Ridge | 243.7 ± 41.1 | 3331.9 ± 332.4 | 237.4 ± 54.7 | 268.6 ± 47.2 |
| RF | 141.3 ± 31.1 | 1495.6 ± 209.0 | 154.9 ± 61.4 | 174.8 ± 23.0 |
| XGBoost | 134.2 ± 20.5 | 1448.5 ± 201.6 | 155.0 ± 66.6 | 163.1 ± 28.7 |
| KNN | 129.9 ± 31.1 | 1358.7 ± 253.4 | 142.2 ± 50.1 | 180.0 ± 18.2 |
| **MLP** | **95.6 ± 12.2** | **1309.5 ± 401.4** | **92.0 ± 26.3** | **138.4 ± 36.1** |
| SVR | 103.6 ± 13.3 | 1326.4 ± 195.9 | 101.5 ± 29.7 | 125.5 ± 20.0 |

**RMSE (nM)**

| Model | E2 | AA | Mel | 5-HT |
|---|---|---|---|---|
| Ridge | 410.5 ± 56.9 | 4492.0 ± 442.1 | 408.1 ± 93.4 | 455.0 ± 93.5 |
| RF | 272.6 ± 54.9 | 2258.3 ± 429.5 | 302.5 ± 102.8 | 363.5 ± 42.4 |
| XGBoost | 272.4 ± 40.2 | 2113.8 ± 341.7 | 290.3 ± 105.2 | 328.2 ± 39.2 |
| KNN | 265.0 ± 74.8 | 2138.9 ± 328.7 | 296.5 ± 70.2 | 375.5 ± 39.3 |
| **MLP** | **182.5 ± 28.8** | **1933.5 ± 559.7** | **181.6 ± 51.8** | **277.3 ± 75.1** |
| SVR | 206.8 ± 25.9 | 2071.7 ± 263.9 | 227.6 ± 56.9 | 277.4 ± 61.3 |

**Overall model performance (avg across analytes)**

| Model | Test $R^2$ Avg | Train $R^2$ Avg | Val $R^2$ Avg | Time (s) |
|---|---|---|---|---|
| Ridge | 0.611 ± 0.063 | 0.728 ± 0.023 | 0.601 ± 0.011 | 1.9 ± 3.2 |
| RF | 0.828 ± 0.043 | 0.984 ± 0.000 | 0.814 ± 0.005 | 512.6 ± 28.6 |
| XGBoost | 0.848 ± 0.041 | 0.994 ± 0.000 | 0.830 ± 0.010 | 111.9 ± 5.7 |
| KNN | 0.828 ± 0.065 | 1.000 ± 0.000 | 0.789 ± 0.017 | 1.4 ± 0.2 |
| **MLP** | **0.907 ± 0.030** | **0.958 ± 0.004** | **0.848 ± 0.012** | **462.4 ± 36.8** |
| SVR | 0.892 ± 0.023 | 0.959 ± 0.001 | 0.850 ± 0.009 | 221.4 ± 8.6 |

**Table S5. Literature Comparison.** Comparison of electrochemical regression studies combined with ML since 2019, showing sensor material, biomarkers, lower concentration levels tested, number of samples, electrochemical methods, ML model, and $R^2$. Biosensors based on a bio-recognition element (e.g., aptamers, enzymes) are not included. Note that reported $R^2$ values may correspond to calibration, cross-validation, or held-out test sets depending on the study; direct comparison should therefore be made with caution.

| **Sensor** | **Analytes** | **Con.** | **#** | **Methods** | **ML** | $R^2$ | **Ref** |
|---|---|---|---|---|---|---|---|
| CNT/g-$C_3N_4$-GCE | Morphine, Methadone, UA | µM | 29 | FFT-SWV | PLSR | 0.96 | [36] |
| MWCNT-ZnO-CuO-microflowers | Urea | mM | >200 | CV | KNN | 0.98 | [37] |
| CB–GO/CP | Tyrosine | µM | 32 | DPV | Ridge regression | 0.98 | [38] |
| NiO/Pt, $Ni(OH)_2$/Au, $Ni(OH)_2$/Pt | Glucose, Lactate | mM | 135 | Chrono-amperometry | BP-NN | 1.00 | [39] |
| $Ni(OH)_2$-GCE | Insulin, Glucose | pM, mM | 168 | CV | Linear regression | 0.98-0.99 | [18] |
| ERGO-GCE | DA, 5-HT, UA, AA | µM | 45 | DPV | ANN | 0.97 | [17] |
| GCE | DA, 5-HT | µM | 72 | DPV | PCA-GPR | 0.78 | [40] |
| **PVDF-CF, P4VP-CF** | **E2, 5-HT, Mel, AA** | **nM** | **450** | **SWV** | **MLP** | **0.95** | **This work** |

Abbreviations: CNT: carbon nanotubes; UA: uric acid; GCE: glassy carbon electrode; FFT-SWV: fast Fourier transform square-wave voltammetry; PLSR: partial least-squares regression; MWCNT: multi-walled carbon nanotube; CB–GO/CP: carbon black–graphene oxide conjugate polymer; DPV: differential pulse voltammetry; BP-NN: back-propagation neural network; ERGO: electrochemically reduced graphene oxide.

**Table S6. Preprocessing/Polymer-Dependent Performance.** Test $R^2$ for models trained separately on PVDF-CF or P4VP-CF data (mean ± std, 5 seeds) for all biomarkers. Three preprocessing methods evaluated: **min-centering** (baseline shift to zero), **polynomial** (linear background subtraction), and **derivative** ($dI/dV$ transformation for peak enhancement). Analysis reveals polymer-specific selectivity: PVDF-CF excels at E2/Mel quantification; P4VP-CF at AA/5-HT quantification. Highlighted combinations show expected optimal polymer-preprocessing pairs per analyte.

| **E2** | **PVDF-mincent.** | **P4VP-mincent.** | **PVDF-poly** | **P4VP-poly** | **PVDF-derivative** | **P4VP-derivative** |
|---|---|---|---|---|---|---|
| Ridge | 0.768 ± 0.205 | 0.707 ± 0.148 | 0.309 ± 1.187 | 0.659 ± 0.160 | **0.808 ± 0.193** | 0.715 ± 0.118 |
| RF | 0.752 ± 0.150 | 0.685 ± 0.114 | 0.616 ± 0.321 | 0.735 ± 0.088 | **0.647 ± 0.291** | 0.764 ± 0.076 |
| XGBoost | 0.796 ± 0.094 | 0.724 ± 0.090 | 0.498 ± 0.468 | 0.757 ± 0.050 | **0.708 ± 0.205** | 0.790 ± 0.067 |
| KNN | 0.630 ± 0.262 | 0.665 ± 0.211 | 0.715 ± 0.146 | 0.743 ± 0.088 | **0.814 ± 0.077** | 0.802 ± 0.052 |
| MLP | 0.852 ± 0.055 | 0.774 ± 0.059 | 0.867 ± 0.068 | 0.790 ± 0.084 | **0.825 ± 0.155** | 0.868 ± 0.042 |
| SVR | 0.771 ± 0.129 | 0.636 ± 0.163 | 0.709 ± 0.169 | 0.778 ± 0.121 | **0.664 ± 0.095** | 0.872 ± 0.027 |
| **Mel** | **PVDF-mincent.** | **P4VP-mincent.** | **PVDF-poly** | **P4VP-poly** | **PVDF-derivative** | **P4VP-derivative** |
| Ridge | 0.771 ± 0.212 | 0.658 ± 0.108 | **0.867 ± 0.076** | 0.686 ± 0.093 | 0.839 ± 0.113 | 0.713 ± 0.116 |
| RF | 0.628 ± 0.112 | 0.619 ± 0.158 | **0.875 ± 0.072** | 0.757 ± 0.076 | 0.837 ± 0.099 | 0.783 ± 0.080 |
| XGBoost | 0.693 ± 0.109 | 0.668 ± 0.142 | **0.861 ± 0.061** | 0.777 ± 0.076 | 0.804 ± 0.118 | 0.815 ± 0.062 |
| KNN | 0.581 ± 0.135 | 0.587 ± 0.225 | **0.843 ± 0.080** | 0.743 ± 0.119 | 0.845 ± 0.067 | 0.758 ± 0.125 |
| MLP | 0.791 ± 0.184 | 0.753 ± 0.094 | **0.960 ± 0.029** | 0.881 ± 0.057 | 0.923 ± 0.075 | 0.892 ± 0.050 |
| SVR | 0.840 ± 0.163 | 0.646 ± 0.142 | **0.894 ± 0.095** | 0.766 ± 0.051 | 0.864 ± 0.128 | 0.806 ± 0.075 |
| **5-HT** | **PVDF-mincent.** | **P4VP-mincent.** | **PVDF-poly** | **P4VP-poly** | **PVDF-derivative** | **P4VP-derivative** |
| Ridge | 0.543 ± 0.185 | 0.575 ± 0.195 | 0.534 ± 0.176 | 0.577 ± 0.180 | 0.617 ± 0.190 | **0.634 ± 0.156** |
| RF | 0.431 ± 0.253 | 0.630 ± 0.201 | 0.429 ± 0.232 | 0.774 ± 0.099 | 0.686 ± 0.176 | **0.724 ± 0.159** |
| XGBoost | 0.523 ± 0.271 | 0.611 ± 0.319 | 0.453 ± 0.275 | 0.778 ± 0.053 | 0.680 ± 0.203 | **0.772 ± 0.133** |
| KNN | 0.348 ± 0.211 | 0.531 ± 0.163 | 0.446 ± 0.174 | 0.650 ± 0.178 | 0.455 ± 0.295 | **0.720 ± 0.154** |
| MLP | 0.718 ± 0.153 | 0.727 ± 0.167 | 0.619 ± 0.191 | 0.807 ± 0.135 | 0.619 ± 0.217 | **0.860 ± 0.126** |
| SVR | 0.673 ± 0.215 | 0.389 ± 0.417 | 0.726 ± 0.202 | 0.717 ± 0.195 | 0.725 ± 0.218 | **0.850 ± 0.070** |
| **AA** | **PVDF-mincent.** | **P4VP-mincent.** | **PVDF-poly** | **P4VP-poly** | **PVDF-derivative** | **P4VP-derivative** |
| Ridge | 0.560 ± 0.269 | 0.421 ± 0.123 | 0.651 ± 0.137 | 0.504 ± 0.099 | 0.685 ± 0.099 | **0.452 ± 0.087** |
| RF | 0.794 ± 0.119 | 0.870 ± 0.053 | 0.726 ± 0.184 | 0.875 ± 0.044 | 0.845 ± 0.079 | **0.831 ± 0.054** |
| XGBoost | 0.811 ± 0.064 | 0.903 ± 0.042 | 0.731 ± 0.109 | 0.871 ± 0.043 | 0.867 ± 0.058 | **0.859 ± 0.036** |
| KNN | 0.690 ± 0.173 | 0.833 ± 0.084 | 0.570 ± 0.263 | 0.823 ± 0.077 | 0.732 ± 0.062 | **0.885 ± 0.023** |
| MLP | 0.792 ± 0.114 | 0.867 ± 0.110 | 0.806 ± 0.081 | 0.890 ± 0.061 | 0.839 ± 0.071 | **0.926 ± 0.020** |
| SVR | 0.744 ± 0.169 | 0.366 ± 0.117 | 0.732 ± 0.087 | 0.680 ± 0.090 | 0.842 ± 0.061 | **0.889 ± 0.061** |
